# Cotranslational Folding Stimulates Programmed Ribosomal Frameshifting in the Alphavirus Structural Polyprotein

**DOI:** 10.1101/790444

**Authors:** Haley R. Harrington, Matthew H. Zimmer, Laura M. Chamness, Veronica Nash, Wesley D. Penn, Thomas F. Miller, Suchetana Mukhopadhyay, Jonathan P. Schlebach

## Abstract

Viruses maximize their genetic coding capacity through a variety of biochemical mechanisms including programmed ribosomal frameshifting (PRF), which facilitates the production of multiple proteins from a single transcript. PRF is typically stimulated by structural elements within the mRNA that generate mechanical tension between the transcript and ribosome. However, in this work we show that the forces generated by the cotranslational folding of the nascent polypeptide chain can also enhance PRF. Using an array of biochemical, cellular, and computational techniques, we first demonstrate that the Sindbis virus structural polyprotein forms two competing topological isomers during biosynthesis at the ribosome-translocon complex. We then show that the formation of one of these topological isomers is linked to PRF. Coarse-grained molecular dynamic simulations reveal that the translocon-mediated membrane integration of a transmembrane domain upstream from the ribosomal slip-site generates a force on the nascent polypeptide chain that scales with observed frameshifting. Together, our results demonstrate that cotranslational folding of this protein generates a tension that stimulates PRF. To our knowledge, this constitutes the first example in which the conformational state of the nascent chain has been linked to PRF. These findings raise the possibility that, in addition to RNA-mediated translational recoding, a variety of cotranslational folding and/ or binding events may also stimulate PRF.

## Introduction

Viruses have evolved numerous mechanisms to exploit the host machinery in order to increase the coding capacity of their highly constrained genomes. There are at least 27 viral genera that utilize programmed ribosomal frameshifting (PRF) in order to produce multiple proteins from a single transcript (https://viralzone.expasy.org/860). PRF is genetically encoded, and minimally requires a portion of the transcript that contains a repetitive “slippery” heptanucleotide sequence (slip-site) followed by a region that forms stimulatory RNA secondary structures (an ensemble of stem loops and/ or pseudoknots).^1,2^ A collision between the translating ribosome and the stimulatory secondary structure increases the kinetic barrier to translocation, which causes the ribosome to dwell on the slip-site.^3-7^ During this pause, the t-RNA that is annealed within the ribosomal P-site (and most often also the t-RNA in the A-site)^8^ begin to sample alternative base pairing interactions that shift the reading frame of the ribosme.^9^ Based on these mechanistic considerations, PRF is typically believed to be mediated at the level of RNA structure. Nevertheless, recent reports have also found that the efficiency of −1PRF can be tuned by a variety of regulatory proteins and/ or miRNA.^10-12^

-1PRF is utilized to temporally and stoichiometrically regulate protein production during viral replication and assembly. For instance, the alphavirus structural proteins are most often produced from a single polyprotein that is cleaved into the capsid (CP), E3, E2, 6K, and E1 proteins (Figure 1A).^13^ The E2 and E1 proteins are membrane glycoproteins that heterodimerize early in the assembly pathway. These dimeric units then form trimeric spike complexes, traffic to the plasma membrane, and initiate viral budding.^14-16^ −1PRF during the translation of the 6K protein gives rise to a secondary form of the polyprotein containing the TransFrame (TF) protein,^13,17^ a known virulence factor,^18-21^ in place of the 6K and E1 proteins (Figure 1B). Because −1PRF precludes the translation of E1, the efficiency of ribosomal frameshifting (1-48% in alphaviruses)^22^ influences the stoichiometric ratio of the E1 and E2 glycoproteins and the net accumulation of spike complexes. Current evidence suggests −1PRF is stimulated by a canonical poly-U slip site and a downstream RNA hairpin.^23^ However, an effort to map the stimulatory RNA structures within alphavirus polyproteins revealed that deletions within the predicted hairpin region are capable of reducing the efficiency of −1PRF, but appear to be insufficient to knock out frameshifting completely.^22^ This observation suggests there may be multiple regulatory mechanisms that mediate −1PRF within the alphavirus structural polyprotein.

**Figure 1.**
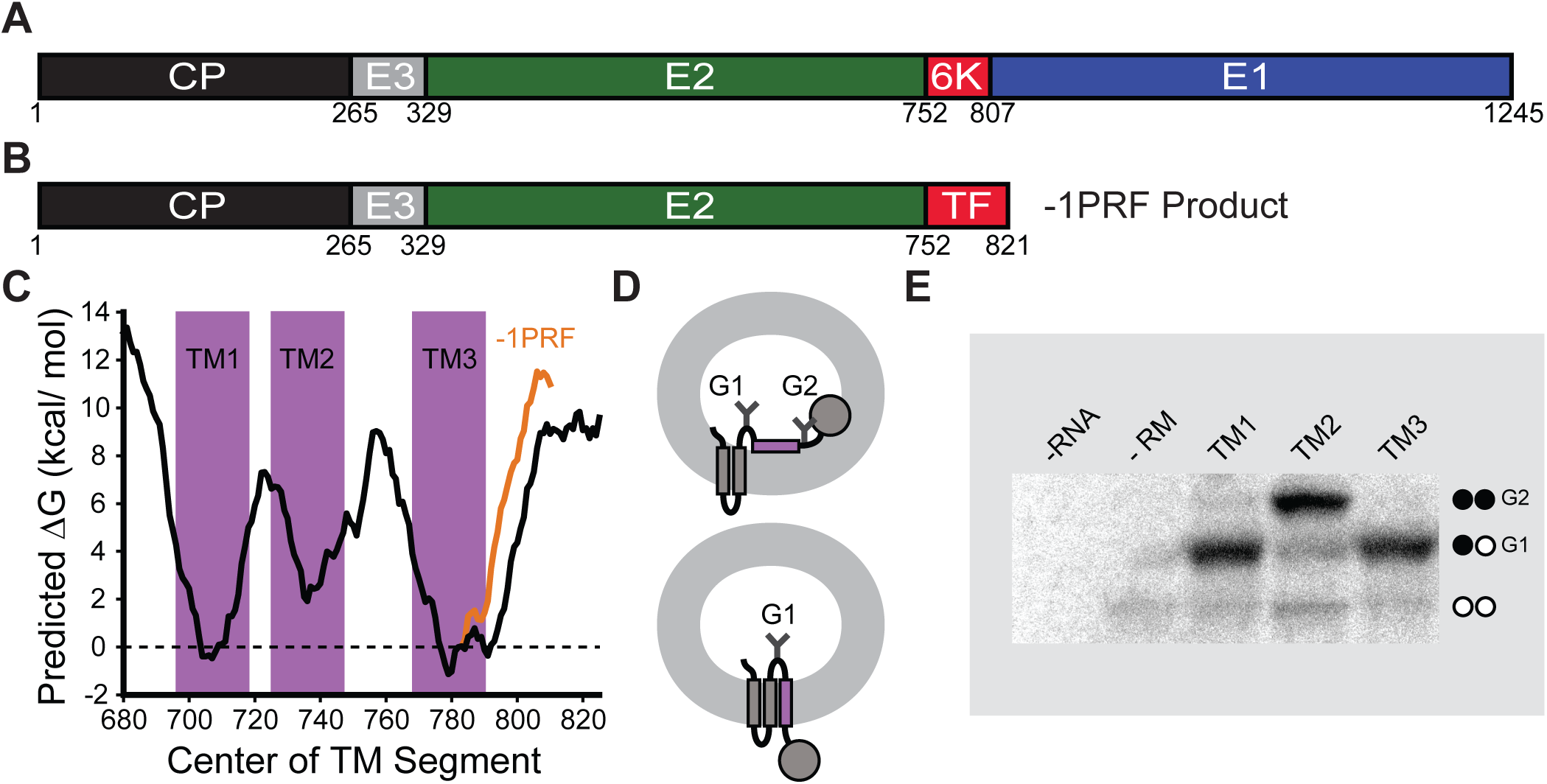
Structure and Topological Properties of the Alphavirus Polyprotein. A) A cartoon depicts the relative size and orientation of proteins within the major form of the alphavirus structural polyprotein. B) A cartoon depicts the relative size and orientation of proteins within the frameshifted form of the alphavirus structural polyprotein. C) A portion of the SINV structural polyprotein spanning the E2, 6K, and E1 proteins was scanned with the ΔG predictor using a 23 residue window.^33^ The predicted free energy difference associated with the cotrantlational membrane integration of every possible 23 residue segment within the major form (black) and frameshifted form (orange) of the structural polyprotein is plotted as a function of the position of its central residue. The position of each predicted TM domain is indicated in purple. D) A cartoon depicts the manner in which the topological preferences of the guest TM domain (purple) influence the glycosylation state of the chimeric Lep protein. The glycosylation machinery is on the interior of the vesicle (the microsomal lumen). Membrane integration of the guest domain results in the production of a singly glycosylated product (bottom). The mis-integration of the guest domain results in the production of a doubly-glycosylated product (top). E) Chimeric Lep constructs bearing putative TM domains from the SINV structural polyprotein were produced by *in vitro* translation in the presence of canine rough microsomes, and analyzed by SDS-PAGE. A representative gel reveals the relative abundance of singly (G1) and doubly (G2) glycosylated translation products for each contruct. Control reactions contaning no RNA (no protein) and no rough microsomes (untargetted protein) are shown for the sake of comparison. These trends were consistently observed across five independent replicates.

-1PRF occurs during synthesis and processing of the nascent alphavirus structural polyprotein at the endoplasmic reticulum (ER) membrane. Following autoproteolytic cleavage of CP in the cytosol, a signal peptide at the N-terminus of the E3 protein directs the nascent polyprotein to the endoplasmic reticulum (ER) lumen where processing of the downstream proteins occurs. Localization of these segments within the lumen is essential to ensure that the E3, E2, and E1 ectodomains form their native disulfides and undergo glycosylation.^15,16,24^ Post-translational modifications are also critical for TF, which must be palmitoylated in order to reach the plasma membrane and incorporate into the viral envelope.^25^ The palmitoylated cysteines in TF are positioned near the edge of a putative transmembrane (TM) domain that is found in both TF and 6K.^25,26^ Though these residues are present in both proteins, they are only palmitoylated in the context of the frameshifted polyprotein.^25^ Considering palmitoylation only occurs on the cytosolic face of cellular membranes,^27^ the distinct modification state of the two forms of the polyprotein is therefore suggestive of an underlying difference in their topologies. In this study, we set out to gain insight into the interplay between −1PRF and the topology of the structural polyprotein. We first mapped the topology of the Sindbis virus (SINV) structural polyprotein. Our results demonstrate that the structural polyprotein forms two topological isomers. The predominant topology features two TM domains upstream of the −1PRF site, and its formation coincides with production of the 6K protein. Alternatively, the minor topology contains an additional TM domain upstream from the −1PRF site that is linked to the production of TF. Using protein engineering in conjunction with coarse-grained molecular dynamics (CGMD) simulations, we demonstrate that the efficiency of −1PRF is dependent upon the force generated by the translocon-mediated membrane integration of the extra TM domain within the minor topomer. Together, our observations highlight novel connections between the cotranslational folding, biosynthesis, and processing of the alphavirus structural polyprotein. Moreover, our findings reveal a novel mechanism that regulates the overall efficiency of −1PRF.

## Results & Discussion

### Topological Properties of the Alphavirus Structural Polyprotein

The current model of the alphavirus structural polyprotein suggests the E2 and 6K proteins each contain two TM domains.^13,28^ However, there are two caveats to this model. First, cryo-EM structures reveal that the E2 protein only contains a single TM domain in the context of the viral envelope.^29,30^ Though it has been speculated that a second TM domain within E2 is somehow extruded from the membrane during processing, the marginal hydrophobicity of this segment also raises the possibility that it may fail to undergo translocon-mediated membrane integration in the first place. Second, the hydrophobic portion of the SINV 6K protein is only 35 residues in length, which is quite short for a segment containing two putative TM domains and a loop. These ambiguous topological signals suggest this portion of the polyprotein is frustrated and could potentially form multiple topological isomers,^31^ as has been suggested for the coronavirus E protein.^32^

To survey the topological preferences of the E2-6K region, we scanned its sequence using a knowledge-based algorithm that predicts the energetics associated with the transfer of polypeptide segments from the translocon to the ER membrane (ΔG predictor).^33^ Energetic predictions suggest only the regions corresponding to the first hydrophobic segments within the E2 (TM1) and 6K (TM3) proteins are sufficiently hydrophobic to undergo robust membrane integration (Δ*G* < 0 kcal/ mol, Fig. 1C). In contrast, the translocon-mediated membrane integration of the second hydrophobic segment within E2 (TM2) is predicted to be inefficient (Fig. 1C). To test these predictions, we measured the translocon-mediated membrane integration of each putative TM domain using a glycosylation-based translocation assay.^34^ Briefly, the sequences of each putative TM domain were cloned into a chimeric leader peptidase (Lep) reporter protein.^34^ Membrane integration of the putative TM segment (purple helix, Fig. 1D) results in the modification of only a single glycosylation site in Lep, whereas the passage of the segment into the lumen results in the modification of two glycosylation sites (Fig. 1D). Chimeric Lep proteins were produced by *in vitro* translation in the presence of canine rough microsomes, which contain native ER membranes and translocons. Consistent with predictions, Lep proteins containing TMs 1 & 3 acquire a single glycosyl modification, which suggests these segments undergo robust translocon-mediated membrane integration (Fig. 1E). In contrast, the translocon-mediated membrane integration of the second putative TM domain of E2 (TM2) is significantly less efficient (Fig. 1E). These observations suggest the E2 and 6K proteins are each likely to contain a single TM domain (TMs 1 & 3, Fig. 2A). It should also be noted that −1PRF only modifies the sequence of the loops downstream from these TM domains (Fig. 1C, orange line), and is therefore unlikely to impact their topological preferences.

**Figure 2.**
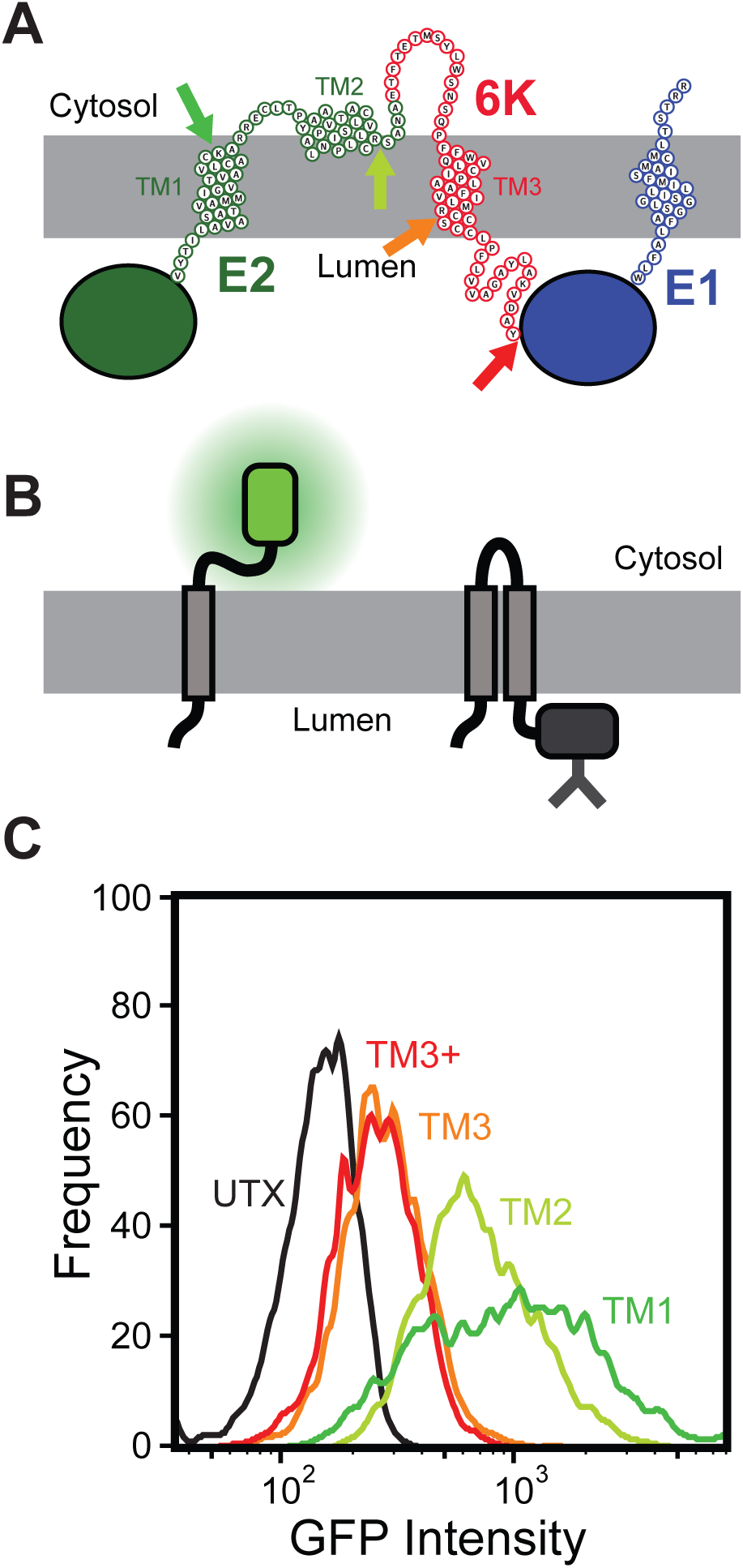
Topological Properties of the Major Form of the SINV Structural Polyprotein. A) A cartoon depicts a putative topological model of most abundant topology of the structural polyprotein that is consistent with computational and biochemical data. The positions at which glycosylatable GFP (gGFP) reporter domains were fused to determine the cellular compartmentalization of the C-terminal portion of the segments corresponding to TMs 1 (green), 2 (yellow), 3 (orange), and the C-terminal edge of the hydrophobic portion of 6K (TM3+, red) are indicated with arrows. B) A cartoon depicts the manner in which the cellular compartmentalization of the gGFP reporter domain alters its fluorescence. Topological signals that direct the gGFP domain into the cytosol will generate a fluorescent gGFP (left), wheras topological signals that direct the gGFP domain into the lumen (right) generate a glycosylated, non-fluorescent fusion domain. C) Reporter constructs bearing a gGFP fusion downstream from TM1 (green), TM2 (yellow), TM3 (orange), and the hydrophobic portion of the 6K protein (TM3+, red) were expressed in HEK293T cells, and cellular fluorescence intensities were analyzed by flow cytometry. A histogram from a representative trial depicts the gGFP intensities assoicated with 3,000 transfected cells expressing each reporter construct at a consistent expression level as judged by the intensity of a bicistronic expression reporter.

Based on the computational and biochemical results in Figure 1, we generated a topological model of the SINV structural polyprotein (Fig. 2A). This model correctly places the E2 and E1 ectodomains within the ER lumen, and places the two palmitoylated cysteine residues in E2 (C716 and C718) within the cytosol.^35,36^ To probe the topological preferences of the SINV polyprotein in the cell, we produced and characterized a series of reporter constructs that begin with the E3 protein and end at the C-terminal edge of each of the three putative TM domains within E2 and 6K (Fig. 2A, Supplemental Fig. 1). Each of these fragments was genetically fused to a C-terminal cassette containing a short GS linker and glycosylatable GFP (gGFP) gene, which contains two glycosylation sites within the core of the eGFP protein.^37^ Topological signals that direct the gGFP protein into the cytosol will produce a fluorescent gGFP, whereas the glycosylation of gGFP within the ER lumen renders the protein non-fluorescent (Fig. 2B). Each construct was then expressed in HEK293T cells, and flow cytometry was used to quantify the fluorescence intensity of the gGFP reporter at a consistent expression level as judged by the intensity of a bicistronic reporter protein (Supplemental Fig. 2). Expression of the reporter constructs containing gGFP downstream from TMs 1 & 2 (see Fig. 2A) generates fluorescent gGFP (Fig. 2C), which suggests the C-termini of these TM domains reside within the cytosol. In contrast, the reporter construct with gGFP downstream from TM3 (after R785) exhibits an attenuated GFP signal at an equivalent expression level (Fig. 2C), which suggests the gGFP fused to the C-terminus of TM3 is projected into the ER lumen. Placement of the gGFP after the full stretch of hydrophobic amino acids in the 6K protein (after Y807, see Fig. 2A) also results in an attenuated gGFP signal (TM3+ in Fig. 2C), which confirms the full stretch of hydrophobic residues near TM3 only spans the membrane once. Thus, results from this cellular reporter assay (Fig. 2C) are consistent with predictions (Fig. 1C), *in vitro* translation data (Fig. 1E), and the model shown in Figure 2A. These observations together confirm that the E1 and 6K proteins each contain a single TM domain in the most abundant form of the polyprotein.

### Link between Topology and −1 Programmed Ribosomal Frameshifting

The topological properties of the structural polyprotein described above have implications for the manner in which it is processed at the ER membrane. Our model suggests the cluster of unmodified cysteines in the 6K protein (C786, C787, C789, and C790) reside at a C-terminal portion of TM3 that is projected into the ER lumen (Fig. 3A) and is therefore inaccessible to palmitoylating enzymes. However, these same residues are palmitoylated in the TF protein,^25^ which suggests the orientation of TM3 must become inverted upon frameshifting in order to expose them to the cytosolic leaflet. Such an inversion could potentially occur as a consequence of the membrane integration of TM2 (Fig. 3B), which exhibits a weak propensity to undergo translocon-mediated membrane integration (Fig. 1E). Furthermore, the efficiency associated with the translocon-mediated membrane integration of TM2 (∼ 20 %, Fig. 1E) is comparable to the frequency of - 1PRF in the SINV polyprotein (∼16%).^22^ Taken together, these observations potentially suggest a connection between the formation of a secondary topomer and −1PRF.

**Figure 3.**
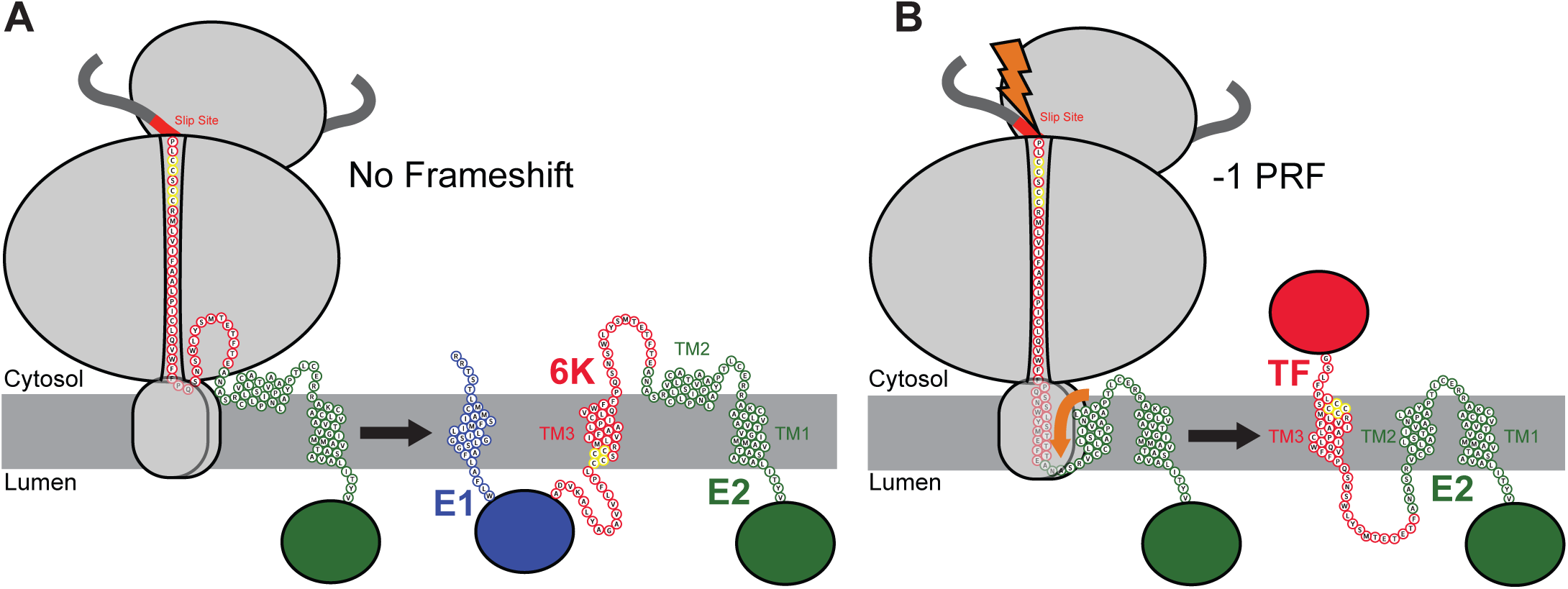
Putative Model for the Interplay Between Translocon Mediated Membrane Integration and Ribosomal Frameshifting. A cartoon depicts the putative manner in which cotranslational folding of the nascent structural polyprotein is linked to −1 programmed ribosomal frameshifting (−1PRF). A) A cartoon depicts the topological properties of the major form of the nascent structural polyprotein. TM2 is too polar to robustly partition into the membrane during translation (left). As a result, the E2 protein only contains a single TM domain in the context of the major form of the structural polyprotein (right). Cysteine residues that are conditionally glycosylated in the frameshifted form of the polyprotein are shown in yellow. B) A cartoon depicts the topological properties of the frameshifted form of the nascent structural polyprotein. TM2 is hydrophobic enough to occaisionally partition into the membrane during translation (left), which imposes a tension on the ribosome that stimulates −1PRF. As a result, the E2 protein contains two TM domains in the context of the frameshifted form of the structural polyprotein (right). Cysteine residues that are conditionally glycosylated in the frameshifted form of the polyprotein are shown in yellow.

Based on these observations, we hypothesize that the translocon-mediated membrane integration of TM2 is mechanistically linked to −1PRF and the translation of the TF protein. Our model suggests mutations that alter the translocon-mediated membrane integration of TM2 should have a direct impact on −1PRF (Fig. 3). To test this hypothesis, we assessed whether mutations that alter the hydrophobicity of TM2 also influence −1PRF. Both energetic predictions and *in vitro* translation measurements suggest the introduction of two non-native leucine residues into TM2 (T738L & S739L, LL mutant) enhances the translocon-mediated membrane integration of TM2 (predicted ΔΔG = −1.7 kcal/ mol, Supplemental Fig. 3). In contrast, the introduction of two glutamate residues into TM2 (V735E & I736E, EE mutant) is predicted to increase its transfer free energy by 3.3 kcal/ mol, which should significantly reduce its membrane integration efficiency (Supplemental Fig. 3A). The effects of the EE substitutions appear to be subtle in the context of the Lep protein (Supplemental Fig. 3B), though this likely reflects the limited dynamic range of this translocation assay.^34,38^ Nevertheless, these results clearly show that the LL and EE mutations alter the translocon-mediated membrane integration of TM2.

To determine whether the cotranslational membrane integration of TM2 impacts translational recoding, we measured the effects of these substitutions on ribosomal frameshifting. PRF is most commonly measured using dual-luciferase reporters, which fuse luciferase domains to the 5’ (renilla luciferase, 0-frame) and 3’ (firefly luciferase, −1 frame) of the gene of interest. The activity of firefly luciferase serves as a reporter for −1PRF and is normalized relative to translational efficiency based on the activity of renilla luciferase. Current versions of these reporter constructs contain self-cleaving 2A segments that release these luciferase domains from the polypeptide of interest.^39^ While the 2A elements are likely to somewhat efficiently release each fusion domain at some point during translation, the introduction of a soluble N-terminal domain could potentially compromise the fidelity of SRP-mediated targeting of the nascent chain to the translocon. To preserve the integrity of the signal peptide, we generated a series of reporter constructs in which translation begins at the native E3 signal peptide and continues until the ribosome reaches a fluorescent mKate fusion domain that is encoded in the −1 reading frame downstream from the PRF site (Supplemental Fig. 4). To control for variations in transfection efficiency at the single-cell level, we included a downstream IRES cassette that drives the bicistronic expression of GFP from the reporter transcript. Reporter constructs encoding TM2 variants of the polyprotein were expressed in HEK293T cells, and cellular mKate intensities were quantitatively compared across cells within a discrete range of IRES-GFP intensities by flow cytometry (Supplemental Fig. 5). The average mKate intensity among cells expressing a reporter construct bearing mutations that knock out the native ribosomal slip-site (UUUUUUA→ GUUCCUA, SSKO) is 79 ± 5% lower than that that among cells expressing the WT form of the reporter (n = 3, Fig. 4A), which confirms that mKate intensities reflect the efficiency of −1PRF. The EE substitutions in TM2 decrease mKate intensity by 61 ± 16% relative to WT (n = 3, Fig. 4A). In contrast, the LL substitutions increase the mKate intensity by 30 ± 11% (n = 3, Fig. 4A). It should be emphasized that each of these reporters contain both the native slip-site and stem loop regions, and that these mutations do not alter their sequences. Thus, these findings demonstrate that −1PRF is sensitive to mutations that impact the membrane integration efficiency of TM2. Together, biochemical evidence suggests that TM2 is inefficiently recognized by the translocon (Fig. 1E), and cellular topology reporters suggest that this segment is most often localized within the cytosol (Fig. 2). Nevertheless, mutagenesis reveals that the propensity of the nascent chain to form a secondary topomer is positively correlated with −1PRF. These results are therefore suggestive of a mechanistic link between topogenesis and −1PRF.

**Figure 4.**
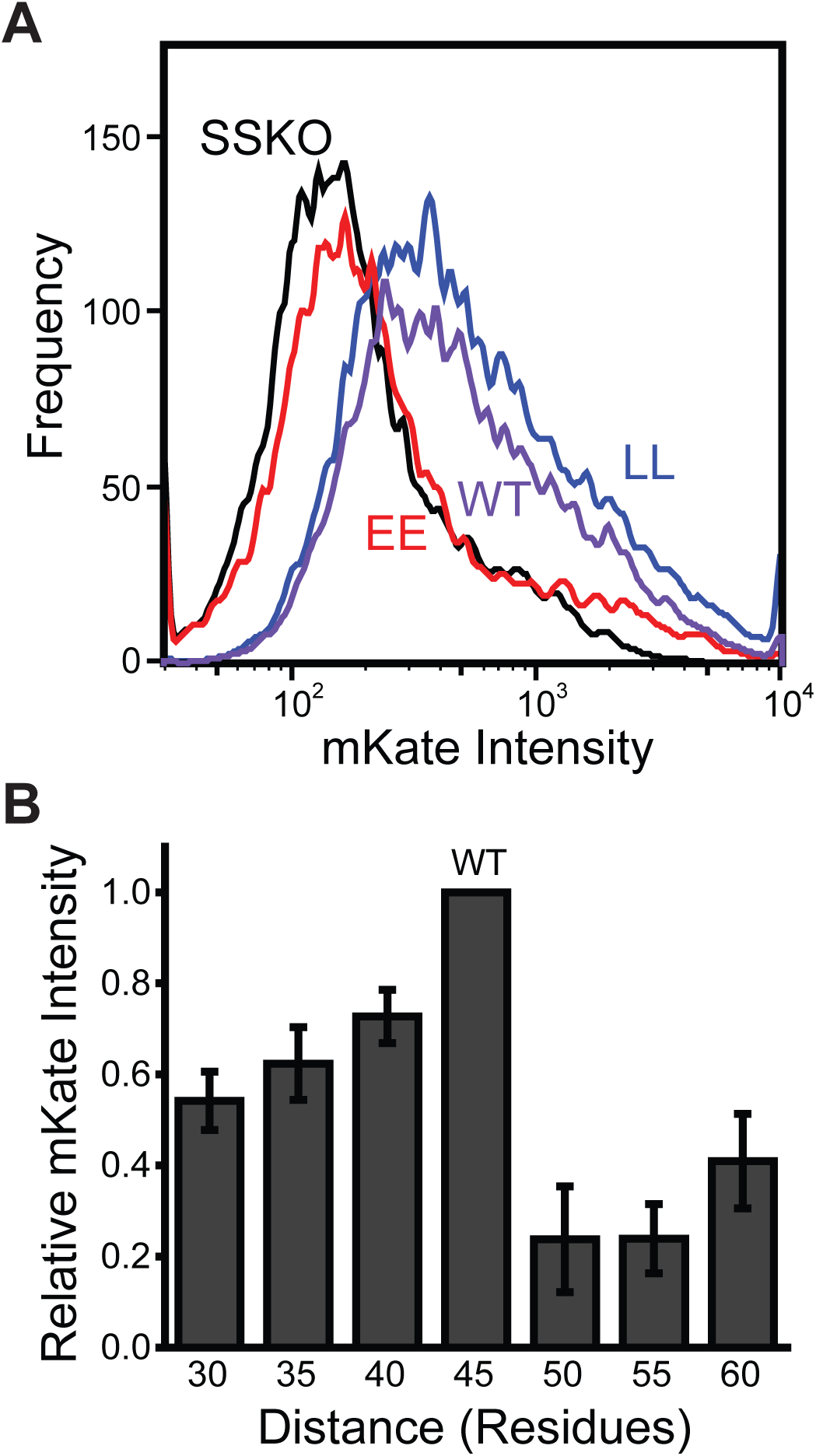
Influence of Sequence Modifications on −1 Programmed Ribosomal Frameshifting. A fragment of the SINV structural polyprotein containing an mKate fused in the −1 reading frame downstream from the ribosomal slip-site was used to compare the effects of sequence modifications on −1PRF levels in HEK293T cells. A) −1PRF reporter constructs containing the WT (black), the EE double mutant (red), and the LL double mutant (blue) TM2 seqeunce were expressed in HEK293T cells, and and cellular fluorescence intensities were analyzed by flow cytometry. A histogram depicts the mKate intensities assoicated with 3,000 cells expressing each reporter construct at a consistent expression level as judged by the intensity of a bicistronic expression reporter. These trends were consistently observed across three independent biological replicates. B) −1PRF reporter constructs containing a series of deletions and G/S linker insertions within the loop between TMs 2 & 3 were expressed in HEK293T cells, and and cellular fluorescence intensities were compared at a consistent expression level by flow cytometry. The mean fluorescence intensities associated with each construct were normalized relative to that of the WT construct, and the average relative fluorescence intensity values are plotted against the number of residues separating the C-terminal residue of TM2 from the slip site. Values represent the average relative fluorescence intensity from three independent biological replicates, and error bars reflect the standard deviation.

### Impact of Nascent Chain Forces on Ribosomal Frameshifting

The apparent link between cotranslational folding and ribosomal frameshifting has implications for the mechanism of −1PRF in the SINV structural polyprotein. The portion of the transcript containing the EE and LL mutations is over 100 nucleotides upstream from the ribosomal slip site, and should therefore not perturb the stimulatory RNA structures that are currently believed to modulate −1PRF.^2,22,23^ These mutations instead alter the portion of the nascent chain that falls just outside of the ribosomal exit tunnel during −1PRF, which suggests the nascent chain itself may stimulate frameshifting. Though it has yet to be implicated in ribosomal frameshifting, the cotranslational membrane integration and/ or folding of the nascent chain is known to generate tension on the ribosome.^41-44^ Furthermore, the C-terminal residue of TM2 is positioned 45 residues upstream of the slip site, which corresponds to a distance that should maximize the tension on the nascent chain at the moment the slip site occupies the ribosomal active site.^41,42^ Previous investigations have demonstrated that the force generated by the membrane integration of the nascent chain is sharply dependent upon this spacing.^41,42^ Therefore, to assess the potential role of this force in ribosomal frameshifting, we generated a set of SINV −1PRF reporter constructs (used in Fig. 4A) containing a series of insertions and deletions that alter the distance between TM2 and the ribosomal slip site (see Supplemental Table 1). Reporter constructs were then expressed in HEK293T cells, and −1PRF reporter intensities were quantitatively compared at a uniform expression level by flow cytometry (Supplemental Fig. 5). A comparison of reporter intensities reveals that −1PRF is maximized at the WT distance of 45 residues (Fig. 4B). In all cases, deletions and insertions that change the distance between TM2 and the slip site significantly reduce the efficiency of −1PRF (Fig. 4B). Moreover, the insertion of a 10-residue G/S linker decreases the intensity of the frameshift reporter by 76 ± 8% (n = 3), which suggests the membrane integration of TM2 is likely to be the primary driver of −1PRF within the SINV structural polyprotein. Nevertheless, the deletion of the region containing the stimulatory RNA hairpin downstream of the slip-site abrogates PRF (Supplemental Fig. 6), which suggests that both the hairpin and TM2 are needed for efficient PRF. Together these findings suggest topological signals within the SINV structural polyprotein generate a mechanical force that stimulates −1PRF.

To further explore the interplay between sequence, topology, and force, we carried out coarse-grained molecular dynamics (CGMD) simulations of the translation and translocon-mediated membrane integration of the nascent structural polyprotein.^44,45^ In these simulations, three-residue segments of the nascent chain are modeled as individual beads with physicochemical properties based on their constituent amino acids. New beads are translated at a rate of 5 amino acids per second, and emerge from the ribosome-translocon complex into an environment with an implicit representation of the bilayer and cytosol (Supplemental Movie 1).^45^ These simulations were previously found to sufficiently recapitulate several aspects of cotranslational membrane protein folding including the formation of topological isomers and the generation of tension on the nascent chain.^44,46,47^ CGMD simulations of SINV polyprotein biosynthesis suggest the nascent chain samples several different topological isomers (Fig. 5A), and that its topological heterogeneity persists after the polyprotein has cleared the translocon. TM2 undergoes translocon-mediated membrane integration (Fig. 5A, right) in only 44 ± 4% of the CGMD trajectories in which TM1 is correctly integrated into the membrane. Consistent with expectations, CGMD simulations suggest that the membrane integration efficiency of TM2 is enhanced by the LL mutations (51 ± 4%) and reduced by the EE mutations (11 ± 3%). This finding provides additional evidence that the topological frustration within this domain (see Fig. 3) arises primarily from its marginal hydrophobicity.

**Figure 5.**
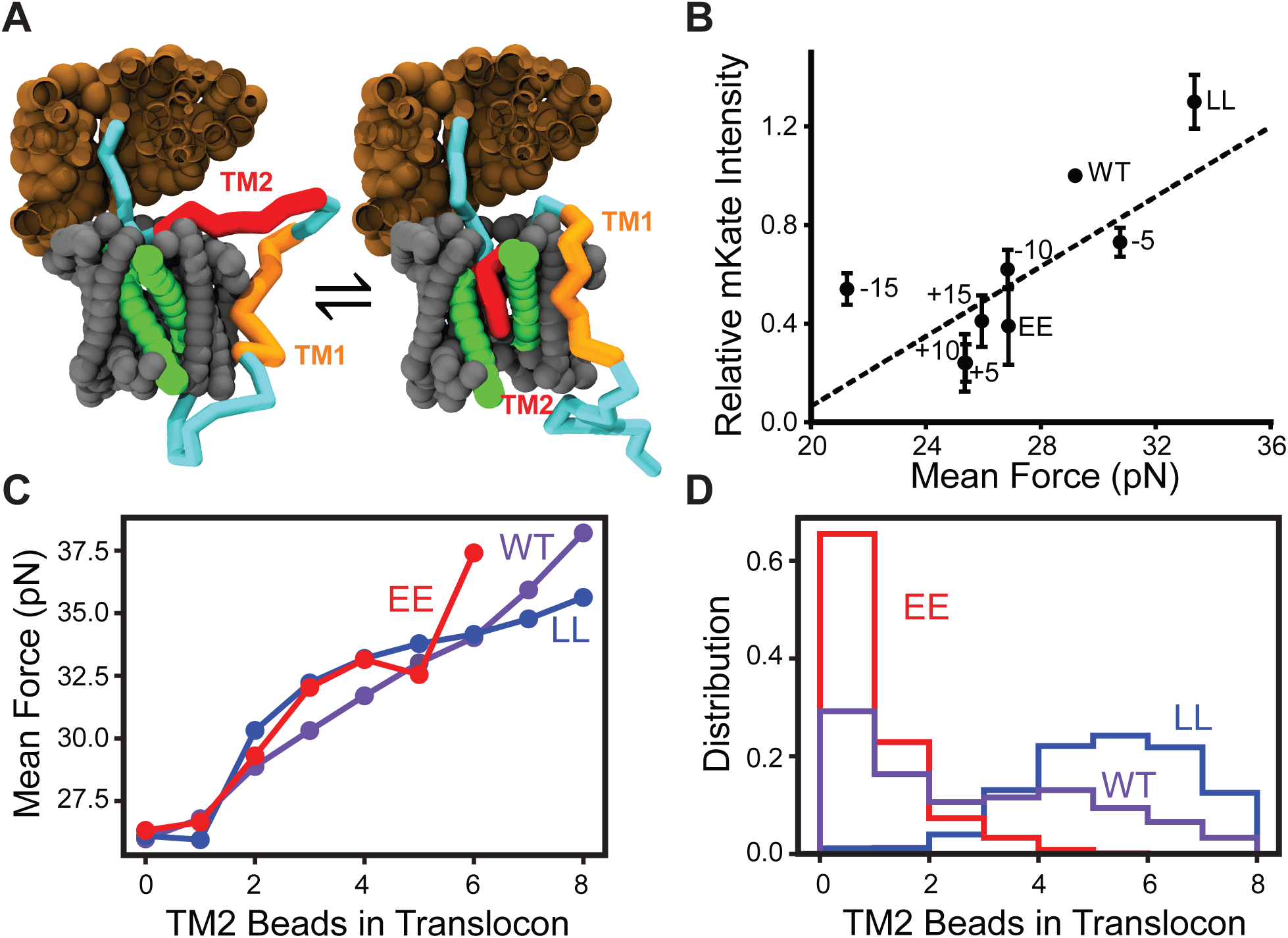
Coarse-Grained Molecular Dynamics Simulations of Polyprotein Biosynthesis. Coarse-grained molecular dynamics (CGMD) simulations were carried out to simulate biosynthesis of a series of SINV structural polyprotein variants, and the pulling force on the nascent chain was calculated while the ribosomal slip site occupies the ribosomal P-site. A) Representative snapshots from CGMD simulations are shown during translation at the slip-site, which is the point during elongation at which pulling forces on the nascent chain are measured. The ribosomal exit tunnel is shown in brown. The translocon is shown in gray and its lateral gate is highlighted in green. The nascent chain is shown in blue, except for the portions that correspond to TMs 1 & 2, which are highlighted in orange and red, respectively. The snapshot on the left depicts a representative trajectory in which TM2 passes through the lateral gate and into the membrane. The snapshot on the right depicts a representative trajectory in which TM2 fails to enter the translocon. B) −1PRF fluorescence reporter (mKate) intensity values for cells expressing a series of modified polyprotein variants were normalized relative to wild-type and plotted against the corresponding mean force measurements calculated from 560 CGMD trajectories. Error bars reflect the standard deviations from three independent biological replicates. The identity of each variant (see Supplemental Table 1) along with a linear fit of the data (dashes) are shown for reference (Pearson’s R = 0.75). C) Pulling force measurements are compared among topological isomers for the WT (purple), LL (blue), and EE (red) polyprotein variants in which the number of TM2 residues (or beads) located within the translocon was found to vary. D) A histogram depicts the number of TM2 residues (or beads) within the translocon among the conformational trajectories sampled during biosynthesis of the WT (purple), LL (blue), and EE (red) variants of the SINV polyprotein.

To evaluate the connection between pulling forces on the nascent chain and frameshifting, we measured the tension on the nascent chain at the point of elongation when the slip site occupies the ribosomal active site (Supplemental Movie 1).^47^ Pulling forces were highest for the LL variant, which averaged 4.2 pN higher than the WT. In contrast, the EE mutations reduce the pulling force on the nascent chain by an average 2.3 pN relative to WT. These results are consistent with the hypothesis that differences in frameshifting arise from the effects of these mutations on the pulling force on the nascent chain. Simulations of polyprotein variants bearing insertions or deletions that alter the distance between TM2 and the ribosomal slip-site indicate that the native distance (45 residues) is nearly optimal for the transmission of pulling force through the nascent chain (Supplemental Table 1), which is consistent with the observed patterns in frameshifting (Fig. 4B). Overall, we find the −1PRF efficiency associated with each polyprotein variant roughly scales with corresponding mean pulling force measurements from CGMD simulations (Pearson’s R = 0.75, Fig. 5B), which strongly suggests that the pulling forces generated by the translocon-mediated membrane integration of the nascent chain stimulate −1PRF.

An analysis of the spectrum of topological states sampled during translation reveals that the magnitude of the pulling force transmitted to the ribosome scales with the number of beads that occupy the translocon (Fig. 5C). This finding suggests pulling forces are generated by the movement of hydrophobic transmembrane segments from the protein-conducting channel of the translocon to the hydrophobic membrane core, as has been established previously.^40,41,47^ This interpretation suggests differences in pulling forces arise from variations in the distribution of topological isomers that form during translation of these variants (Fig. 5D). The LL mutant predominately samples conformations where the majority of TM2 beads are in the translocon (Fig. 5A, right), whereas the EE mutant almost exclusively adopts conformations in which the majority of TM2 beads fall outside of the translocon and within the cytosol (Fig. 5A, left). As passage through the translocon is a prerequisite for membrane integration, the relationship between pulling forces and residence of the nascent chain within the translocon is consistent with our model for structural polyprotein biogenesis (Fig. 3) and confirms that the translocon-mediated membrane integration of TM2 stimulates −1PRF.

### Survey of Frameshifting Elements among Alphavirus Structural Polyproteins

Our model suggests that the hydrophobicity of TM2 and its distance from the slip site are the key determinants of the −1PRF efficiency within the SINV structural polyprotein. To assess whether this mechanism is likely to be operative within other alphaviruses, we surveyed six related structural polyproteins for similar sequence elements. Sequence scans carried out with the ΔG predictor reveal that each form of the alphavirus structural polyprotein contains a marginally hydrophobic TM domain upstream from the ribosomal slip site. Predicted transfer free energies associated with the translocon-mediated membrane integration of these putative TM domains range from +1.4 to +2.7 kcal/ mol (Table 1), which suggests the translocon-mediated membrane integration of these segments is likely to be inefficient. Consistent with this notion, CGMD simulations of the translation of these polyproteins indicate that the membrane integration efficiency of these segments ranges from 33-64% (Table 1). Furthermore, these marginally hydrophobic TM domains reside 44-52 residues upstream of their corresponding −1PRF sites (Table 1), which suggests that the tension generated by their translocon-mediated membrane integration is likely to be propagated back to the slip site.^40,41^ Force measurements derived from CGMD simulations of polyprotein synthesis suggest that the tension in the nascent chain when the slip-site occupies the ribosome is comparable to or greater than the tension generated during translation of the SINV variants characterized herein (Table 1). Taken together, these findings suggest that this −1PRF mechanism is likely to be conserved across the alphavirus genus. Additional investigations are needed to determine how nascent chain- and RNA-mediated −1PRF mechanisms are balanced against one another, and how this mechanistic diversity ultimately influences viral evolution.

**Table 1.** Comparison of the Topological Properties of Alphavirus Structural Polyproteins.

| Strain | Accession Number | Predicted $\Delta G_{app}$ of TM2 (kcal/ mol)* | Membrane Integration Efficiency of TM2** | Distance from TM2 to Slip Site (Residues) | Mean Force (pN)*** |
| --- | --- | --- | --- | --- | --- |
| Sindbis Virus | NC_001547 | 1.9 | 0.44 | 45 | 29.2 |
| Eastern Equine Encephalitis Virus | NC_003899 | 2.4 | 0.51 | 45 | 31.0 |
| Middleburg Virus | EF536323 | 1.6 | 0.33 | 52 | 34.0 |
| Sleeping Disease Virus | NC_003433 | 2.7 | 0.33 | 44 | 25.7 |
| Southern Elephant Seal Virus | HM147990 | 2.1 | 0.34 | 47 | 25.1 |
| Semiliki Forest Virus | NC_003215 | 2.1 | 0.64 | 49 | 33.3 |
| Venezuelan Equine Encephalitis Virus | NC_001449 | 1.4 | 0.33 | 44 | 28.6 |

\* Values reflect the minimum $\Delta G_{app}$ value as determined from a sequence scan of the full-length structural polyprotein using the $\Delta G$ predictor. (33)
\*\* Values are derived from CGMD, and reflect the percentage of trajectories in which TM2 was found to adopt a transmembrane orientation.
\*\*\* Values are derived from CGMD, and reflect the average force on the nascent chain while the ribosomal slip site occupies the ribosomal P-site.

## Conclusions

Using an array of biochemical, cellular, and computational methods we show that the nascent SINV structural polyprotein forms a spectrum of topological intermediates during biosynthesis, and that −1PRF is primarily driven by the translocon-mediated membrane integration of a marginally hydrophobic TM domain within the E2 protein. We also provide evidence to suggest this mechanism is likely to be conserved across the alphavirus genus. To date, the mechanistic basis of −1PRF has been generally attributed to the kinetic effects of mechanochemical forces that are generated by structural elements within the mRNA. Indeed, we do find that PRF in the SINV structural polyprotein is dependent upon the RNA stem loop downstream of the slip-site (Supplemental Fig. 6). Nevertheless, it is clear that the forces generated by the translocon-mediated membrane integration of TM2 dramatically enhance the frameshifting efficiency. To our knowledge, the findings reported herein constitute the first instance in which forces generated by conformational transitions in the nascent polypeptide chain have been implicated in the efficiency of PRF. Though additional investigations are needed to elucidate how pulling forces in the nascent chain physically stimulate −1PRF, a causative role for tension in both the transcript and nascent chain seems plausible given that ribosomal frameshifting fundamentally arises from the movement of the tRNA with respect to the mRNA. It seems likely that cotranslational folding is one of many regulators, which include both host and viral effectors, that tune the net efficiency of PRF. This creates the potential for mechanistic diversity that could provide an evolutionary benefit for alphaviruses, as −1PRF is rendered tunable by either downstream or upstream mutations that impact the stability of the mRNA hairpin or the conformational ensemble of the nascent chain, respectively. This flexibility could also potentially provide the virus with a means of maintaining desired −1PRF levels in the presence host factors that globally regulate −1PRF through mRNA interactions.^12^

It should be noted that the implications of these findings potentially extend beyond the realm of viral proteins. A wide variety of molecular transitions have been found to generate tension within the nascent chain including the folding of soluble domains near the ribosomal exit tunnel^43^ and the translocon-mediated membrane integration of nascent TM domains.^40,41^ These observations suggest the tension in the nascent chain should fluctuate as the structural features emerge from the ribosome (Fig. 6A), which may therefore provide the ribosome with a readout for the progress of cotranslational folding. In the case of the SINV structural polyprotein, the topological frustration within the nascent chain leads to the production of two competing topomers that generate distinct pulling forces on the ribosome in a manner that ultimately impacts the fidelity and processivity of translation (Fig. 6 B & C). Additional investigations are needed to explore the potential relevance of this type of cotranslational feedback to protein homeostasis. Indeed, interactions between the nascent chain and molecular chaperones are known to ratchet polypeptides across the membrane,^48,49^ and may therefore contribute to pulling forces. This could potentially account for the fact that the deletion of components of the ribosome-associated chaperone complex has been found to attenuate −1PRF in yeast.^50^ Future investigations are needed to evaluate the full range of −1PRF effectors and how these are potentially exploited for regulatory purposes.

**Figure 6.**
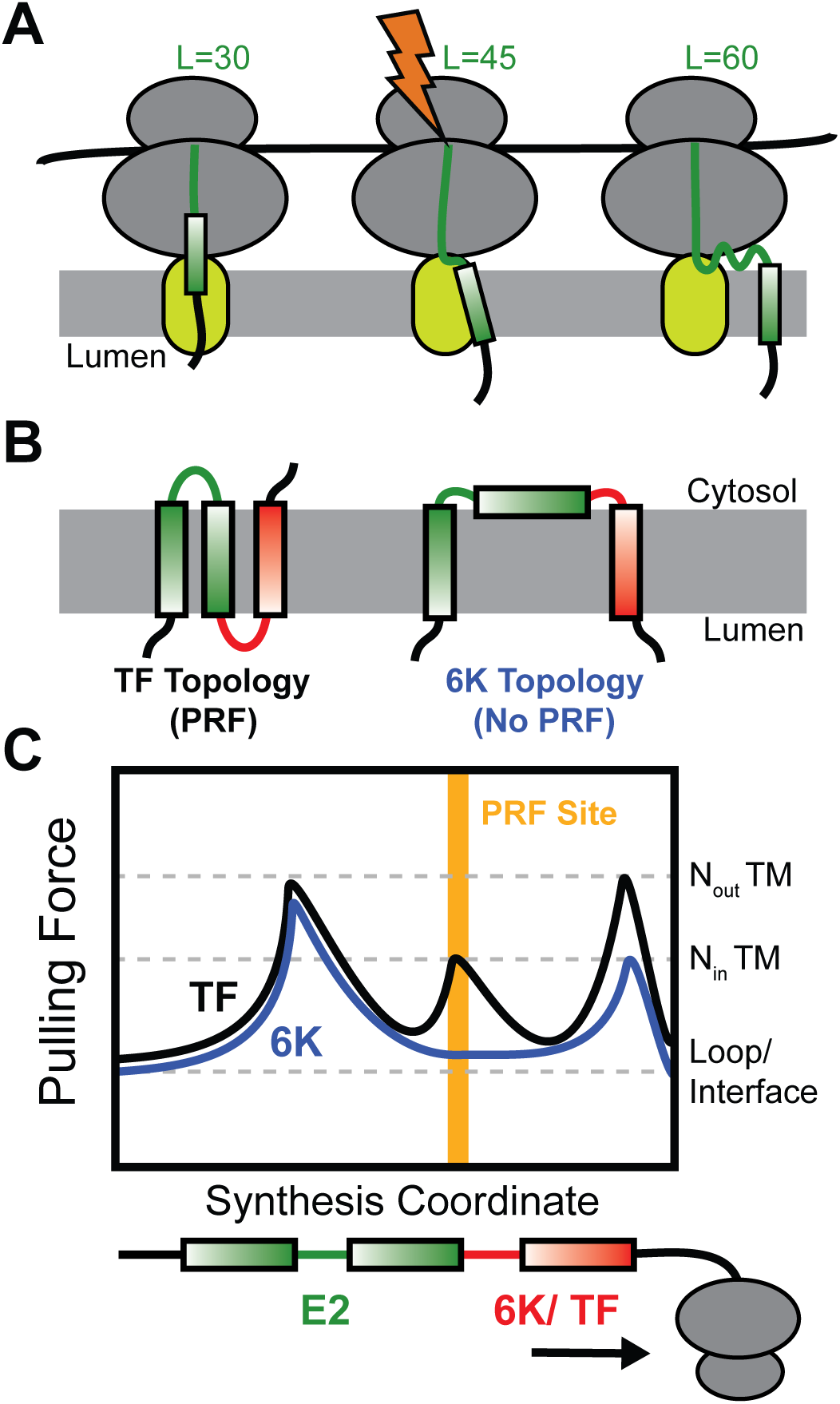
Interplay between Topology, Pulling Force, and Programmed Ribosomal Frameshifting. Cartoons depict the manner in which the translocon-mediated membrane integration of the nascent chain generates a fluctuation in pulling force that triggers PRF during synthesis of the SINV structural polyprotein. A) The pulling force generated by the translocon-mediated membrane integration of each TM domain generates a pulling force on the nascent chain that is maximized during the conjugation of the amino acid that lies ∼45 residues upstream of the C-terminal residue of the TM domain. B) The translocon-mediated membrane integration of TM2 is marginally efficient, which results in the formation of two topologies during translation of the SINV polyprotein. TM2 most often fails to undergo translocon-mediated membrane integration, which results in the formation of a topology featuring only two TM domains (TMs 1 & 3) in the form of the polyprotein containing the 6K protein. However, the translocon-mediated membrane integration of TM2 generates an alternative topology in the frameshifted form of the polyprotein containing the TF protein. C) A hypothetical plot of the fluctuations in the nascent chain pulling force during the translation of the two forms of the SINV structural polyprotein is shown. The translocon-mediated membrane integration of TM2 generates an extra pulling force on the nascent chain while the slip-site occupies the ribosomal P-site, which stimulates −1PRF.

## Methods

### Computational Predictions of Topological Energetics

The energetics associated with the translocon-mediated membrane integration of the nascent structural polyprotein were carried out using the ΔG predictor (http://dgpred.cbr.su.se/).^34^ These predictions are generated using a window scan function that sums depth-dependent free energies associated with the transfer of amino acids from the translocon to the ER membrane.^33^ Full sequence scans of varying window size were used to compare the predicted transfer free energies associated with each segment within the polyprotein in order to identify the segments that are most likely to undergo translocon-mediated membrane integration.

### Plasmid Preparation and Mutagenesis

Chimeric Lep genes were generated in the context of a pGEM-based Lep expression vector^34^ that was kindly provided by the laboratory of Gunnar von Heijne. Putative TM domains of interest were introduced into the H-segment position of this Lep construct using Gibson assembly. To probe the topology of and ribosomal frameshifting within the SINV structural polyprotein, a portion of the polyprotein containing the E3, E2, and 6K/ TF proteins was first introduced downstream from the cytomegalovirus (CMV) promoter within a pcDNA5 vector using Gibson assembly. To produce a series of reporter constructs for polyprotein topology, Gibson assembly was then used to replace the portion of the polyprotein gene downstream from each putative TM domain with a genetic cassette containing a 10-residue G/S linker, a glycosylatable eGFP gene,^37^ an internal ribosomal entry site (IRES), and an mKate gene, respectively (Supplemental Fig. 1).

To produce a series of reporter constructs for ribosomal frameshifting, Gibson assembly was used to replace the portion of the polyprotein gene that falls 100 basepairs downstream from the ribosomal slips site in the 6K gene with a cassette containing an mKate gene in the −1 reading frame followed by an IRES and a GFP gene (Supplemental Fig. 4). The frameshift reporter (mKate) was fused 100 nucleotides downstream from the slip site in order to avoid disrupting the stimulatory RNA hairpin downstream from the slip sites.^22,23^ In addition to avoiding potential issues related to the impact of fusion domains on SRP-targeting of the nascent chain, this design also avoids recently described artefacts associated with previous generations of the dual-luciferase reporter system in two ways.^39^ First, the transcripts of our fluorescent expression reporter (eGFP) does not contain any cryptic splice sites. Second, the fluorescent −1PRF reporter protein (mKate) is liberated from the mutated portion of the nascent polypeptide through a native proteolytic cleavage site between the E2 and 6K/TF protein. Thus, mutations within TM2 should not impact the stability and/ or turnover of mKate. An IRES-eGFP cassette was also incorporated into the downstream portion of the reporter transcript in order to facilitate comparisons of reporter intensities across cells with uniform expression level. Site directed mutagenesis was used to introduce mutations into these constructs. Insertions and deletions were introduced using In-Fusion cloning (Takara, Mountain View, CA).

### In vitro Translation of Chimeric Lep Proteins

Chimeric Lep proteins were generated by *in vitro* translation as was described previously.^51^ Briefly, mRNA for each chimeric Lep protein was produced from plasmids using the RiboMAX RNA production system in accordance with the manufacturer’s instructions (Promega, Madison, WI). Lep proteins were then produced from mRNA by *in vitro* translation using rabbit reticulocyte lysate (Promega, Madison, WI) supplemented with canine pancreatic rough microsomes (tRNA probes, College Station, TX) and EasyTag ^35^S methionine (PerkinElmer, Waltham, MA). *In vitro* translation reactions were then diluted 1:4 into 1X SDS PAGE sample buffer and separated using a 12% SDS PAGE gel. To image radioactive translation products, PAGE gels were then dried, exposed to a phosphorimaging screen, and imaged using a Typhoon Imager (GE Healthcare, New York, NY). Image J software was then employed to process the data quantify the glycosylation state of each construct by densitometry.

### Expression of Fluorescent Reporter Constructs

Flow cytometry was used to compare the fluorescence intensity profiles of HEK293T cells expressing topology and frameshifting reporter constructs. Briefly, HEK293T cells were grown in DMEM (Gibco, Grand Island, NY) containing 10% fetal bovine serum (Corning, Corning, NY) and a penicillin/ streptomycin antibiotic supplement (Gibco, Grand Island, NY) in an incubator containing 5% CO2 at 37° C. Reporter constructs were transiently expressed using lipofectamine 3000 (Invitrogen, Carlsbad CA) in accordance with the manufacturer’s instructions. Cells were harvested two days post-transfection and analyzed using a BD LSRII flow cytometer (BD Biosciences, Franklin Lakes, NJ). Cellular fluorescence profiles were analyzed using FlowJo software (Treestar, Ashland, OR). To compare cellular reporter intensities among cells with a uniform expression level, analysis of reporter intensity levels was restricted to cells that fell within a defined, uniform range of IRES-mKate or IRES-GFP expression reporters intensities. An example of the hierarchical gating strategy employed herein is shown in Supplemental Figs. 2 & 5.

### Course Grained Simulations of Polyprotein Translation

Coarse-grained molecular dynamics (CGMD) simulations are based on a previously developed and tested approach.^44,45^ Briefly, simulations are carried out in the context of a coarsened representation of the ribosome exit tunnel, Sec translocon, and nascent chain. The nascent chain is composed of beads that each represent three amino acids, and new beads are sequentially added to the nascent chain in order to explicitly simulate translation. Translation occurs at a rate of 5 aa/s, which mimics the rate of translation by eukaryotic ribosomes. Each bead interacts with the translocon, ribosome, and other beads in a manner that is dependent on the hydrophobicity and charge of its composite amino acids. Interactions with the solvent and lipid bilayer are modelled implicitly. The ribosome and Sec translocon are fixed in place, with the exception of the lateral gate of the translocon, which stochastically switches between the open and closed conformations in a manner dependent on the free energy difference between the two configurations.

The geometries of the ribosome and translocon are based on cryo-EM structures.^52^ Residue-specific interactions have been parameterized based on over 200 µs of simulations with the MARTINI forcefield. Fitting is performed using a Bayesian uncertainty quantification framework.^53^ This approach represents an update relative previous published methodology, and the new parameters utilized herein are included in Supplemental Table 2. All other parameters necessary to describe the system are available in previous published work.^45^ Integration is performed using overdamped Langevin dynamics with a diffusion coefficient of 253 nm^2^/s and a time step of 300 ns. Despite the significant simplifications involved in this model, the CGMD model has proven accurate in capturing the integration probabilities, topology distributions, and forces experienced in previous studies.^44,45,47^

In order to obtain the distribution of topologies for various polyprotein mutants, the translation and integration of each sequence was simulated 560 times. In order to reduce computational cost, simulations only included the first three TMDs of the alphavirus polyprotein. In order to focus on the topological preferences of TM2, restraints were applied to enforce that TM1 adopts its native topological orientation. Pulling force measurements were performed by pausing translation when the −1PRF site resides within the ribosomal peptidyl transfer center. During this pause, the final bead was fixed in place and the force on the bead exerted by the nascent chain was measured along the translocon channel axis. Due to the truncation of the exit tunnel in our model, the final bead corresponds to the amino acids 27 residues N-terminal of the −1PRF site. Translation is paused for 3 seconds, which is equivalent to the time it would take to translate 5 beads. This relatively short time window ensures that the distribution of polyprotein topologies is not affected by the pause. During this window, pulling forces were measured at a rate of 333 frames per second. In order to sample a wide range of topologies and conformations, each mutant was independently simulated 560 times. This protocol is analogous to force measurements performed in previous work, with exception of a shortened pausing duration.^47^

## Acknowledgments

We thank Renuka Kudva and Gunnar von Heijne for scientific input. We thank David Giedroc and Jonathan Dinman for editorial input. We thank Christiane Hassel and the Indiana University Bloomington Flow Cytometry Core Facility for their experimental support. This research was supported in part by grant from the National Institute of Allergy and Infectious Diseases (NIAID) to J.P.S. and S. M. (R21AI142383) as well as a grant from the National Institute of General Medical Sciences to T. F. M. (R01GM125063).

## Author Contributions

J.P.S, S.M., and T.F.M. III designed the experiments. H.R.H., L.M.C., W.D.P, and V.N. produced the genetic constructs and carried out the biochemical and cellular experiments. M.H.Z. carried out and analyzed CGMD simulations. J.P.S. wrote the manuscript with editorial input from S.M. and T.F.M. III.

## Data Availability

The datasets described herein are available from the corresponding authors upon request. Accession numbers are provided in Table 1.

## Code Availability

All code relating to the CGMD simulations detailed herein have been previously detailed in references 39 and 41, and are freely available upon request.

