## Supplemental Materials for "Cotranslational Folding Stimulates Programmed Ribosomal Frameshifting in the Alphavirus Structural Polyprotein"

*This File Includes:*

Supplemental Figure 1
Supplemental Figure 2
Supplemental Figure 3
Supplemental Figure 4
Supplemental Figure 5
Supplemental Figure 6
Supplemental Table 1
Supplemental Table 2

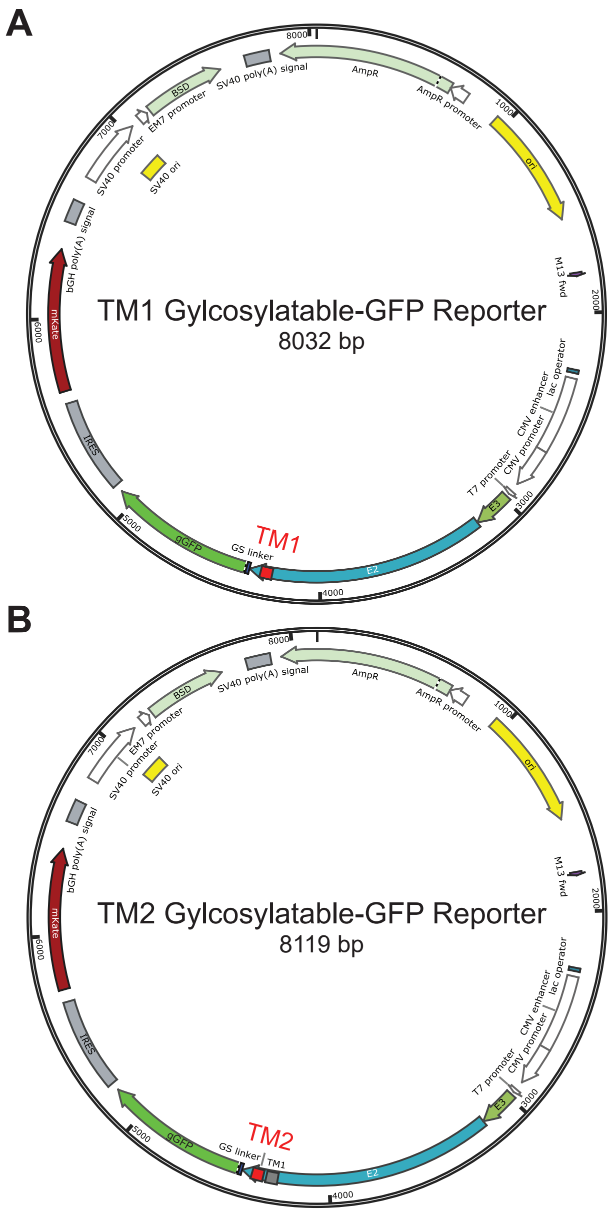


**Supplemental Figure 1. Representative Maps of Topology Reporter Constructs.** Plasmid maps depict the sequence elements cointained within two representative glycosylatable-GFP (gGFP) based topology reporter proteins. Each reporter protein consists of a fragment of the Sindbis structural polyprotein begining with the N-terminal signal peptide in the E3 protein and ending with a putative TM domain of interest, which are expressed from a CMV promoter. A cassette containing a short Gly-Ser linker and gGFP gene is fused downstream of the TM domain of interest. A) A map of the reporter construct for TM1 is shown. B) A map of the reporter construct for TM2 is shown.This reporter is identical to the TM1 reporter, except that the E2 protein is extended through the second TM2 domain.

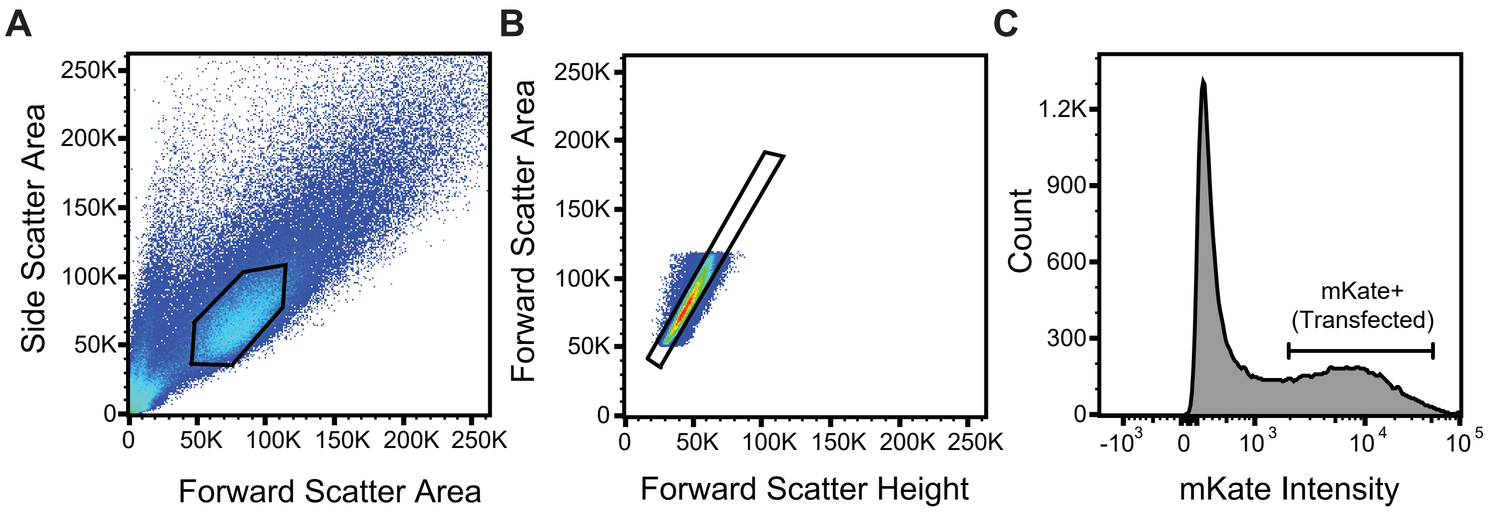


**Supplemental Figure 2. Gating Strategy for the Analsysis Transfected Cells Expressing Topology Reporter Constructs.** The heirarchical gating strategy used to select and analyze positively-transfected single cells is shown for a representative sample of HEK293T cells expressing the WT form of the -1PRF construct (see Figure 2).A) Intact cells were first selected based on the ratio of side scatter to forward scatter intensity. Selected cells are indicated within the black hexagon. B) Single cells were then selected from the population of intact cells based on the ratio of intensities associated with the forward scatter area and height. Selected cells are indicated within the black rectangle. C) Positively transfected cells were then selected for analysis from the population of single cells based on their characteristic bicistronic expression of the mKate fluorophore. Positively-transfected cells selected for analysis of the glycosylatable-GFP reporter intensities are indicated within the brackets. Gates were uniformly applied to all sampled in the analysis to ensure reporter intensities were compared at uniform expression levels.

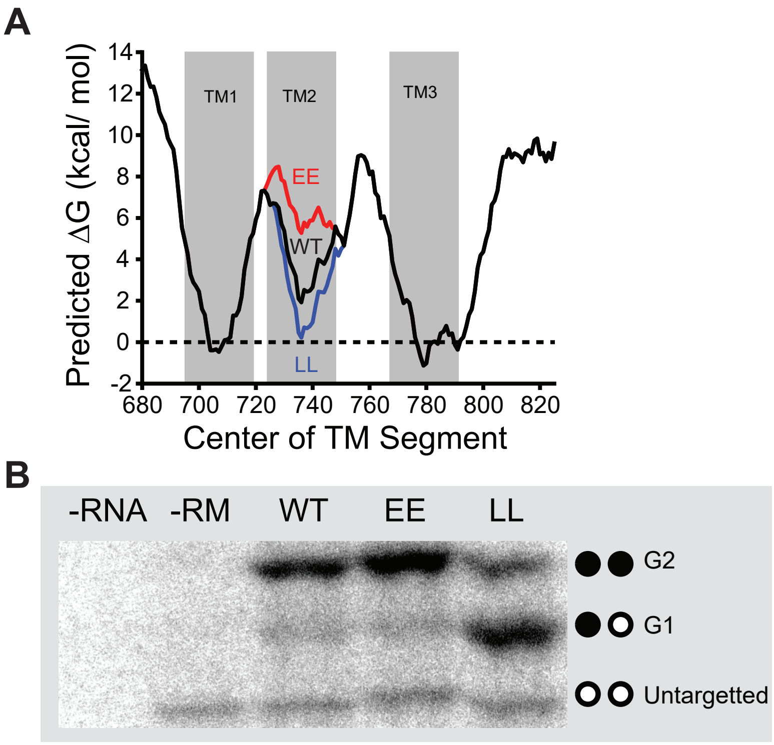


**Supplemental Figure 3. Influence of Mutations on the Translocon-Mediated Membrane Integration of TM2.** A) A portion of the Sindbis structural polyprotein spanning the E2, 6K, and E1 proteins was scanned with the ΔG predictor using a 23 residue window.[20] The predicted free energy difference associated with the cotrantlational membrane integration of every possible 23 residue segment within the structural polyprotein is plotted as a function of the position of its central residue. Transfer free energies associated with the WT form of the polyprotein (black) are shown in relation to those of the V735E/ I736E (EE, red) and T738L/ S739L (LL, blue) double mutants. The position of each predicted TM domain is indicated in gray. B) The translocon-mediated membrane integration of the WT, EE, and LL forms of TM2 were assessed using chimeric leader peptidase (Lep) reporter proteins. Lep constructs bearing the WT, EE, and LL versions of TM2 were produced by *in vitro* translation in the presence of canine rough microsomes and separated by SDS PAGE. A representative gel is shown above, and the position of the untargetted (unglycosylated), singly glycosylated (membrane integrated TM), and doubly glycosylated (lumenal TM) bands are indicated for reference. Negative controls contaning no RNA (no protein) and no rough microsomes (untargetted protein) are shown for reference.

**A**

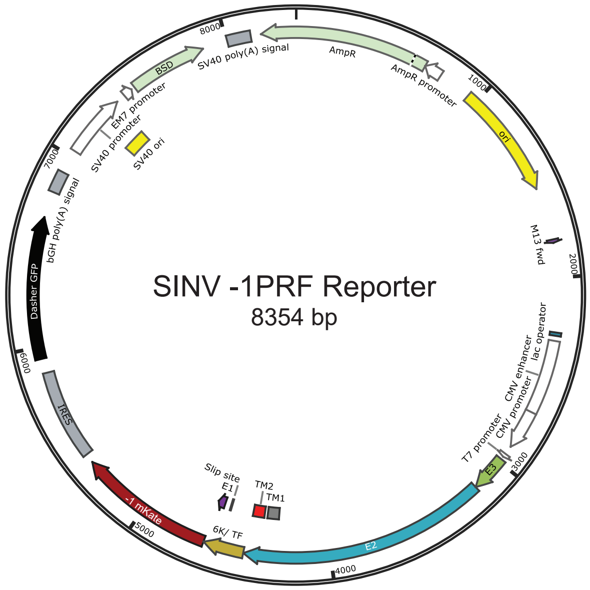


**B**

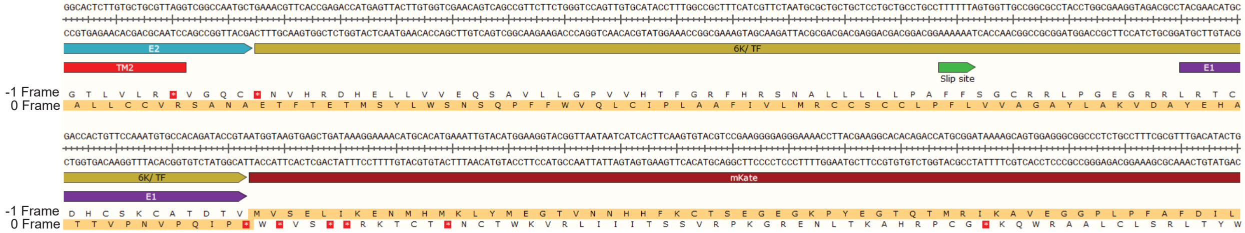


**Supplemental Figure 4. Map of the Sindbis Structural Polyprotein -1PRF Reporter Construct.** Plasmid maps depict the sequence elements cointained within the Sindbis virus structural polyprotein -1PRF reporter protein. A) The -1PRF reporter protein consist of a fragment of the Sindbis structural polyprotein begining with the N-terminal signal peptide in the E3 protein and ending with the N-terminal portion of E1, which is expressed from a CMV promoter. A fluorescent mKate gene is fused to the 3' of this segment in the -1 reading frame, and is not translated in the 0 reading frame B) The sequence of the reporter protein from the end of E2 (blue) through the beginning of mKate (red) is shown for reference. Translation in the 0-frame terminates before the mKate protein. Alternatively, shifting into the -1 reading frame within the ribosomal slip site (green) results in the translation of the fluorescent mKate protein.

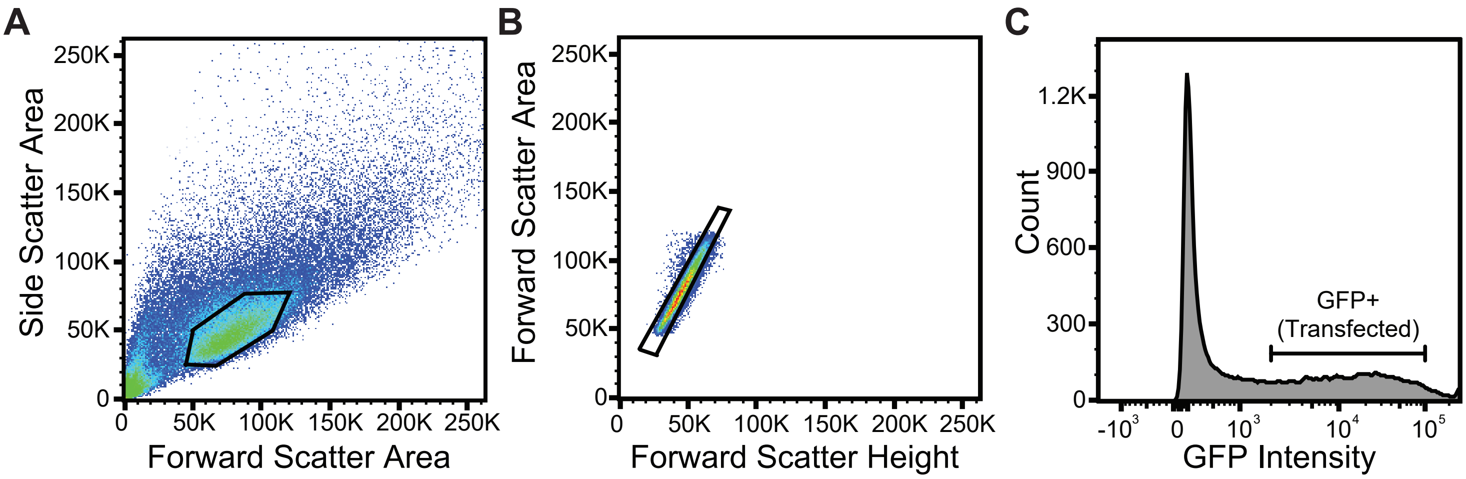


**Supplemental Figure 5. Gating Strategy for the Analsysis Transfected Cells Expressing PRF Reporter Constructs.** The heirarchical gating strategy used to select and analyze positively-transfected single cells is shown for a representative sample of HEK293T cells expressing the WT form of the -1PRF construct (see Figure 4).A) Intact cells were first selected based on the ratio of side scatter to forward scatter intensity. Selected cells are indicated within the black hexagon. B) Single cells were then selected from the population of intact cells based on the ratio of intensities associated with the forward scatter area and height. Selected cells are indicated within the black rectangle. C) Positively transfected cells were then selected for analysis from the population of single cells based on their characteristic bicistronic expression of the GFP fluorophore. Positively-transfected cells selected for analysis of mKate -1PRF reporter intensities are indicated within the brackets. Gates were uniformly applied to all sampled in the analysis to ensure reporter intensities were compared at uniform expression levels.

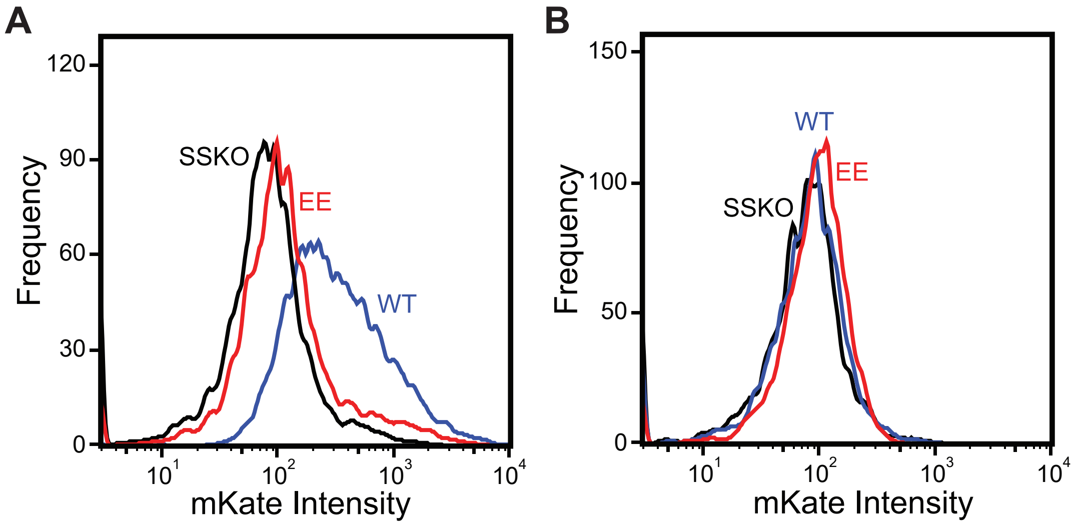


**Supplemental Figure 6. PRF Requires Downstream RNA Structure.** To determine whether the translocon- mediated membrane integration of TM2 is sufficient to independently trigger PRF, we compared the intensity of the fluorescent -1PRF reporter (mKate) generated during the expression of PRF reporter constructs that either (A) contain or (B) lack the downstream RNA hairpin. A) A histogram depicts the distribution of mKate intensities for cells expressing PRF reporter constructs containing the upstream nascent chain with either the WT TM2 (blue) or a mutant form of TM2 bearing to non-native glutamate residues (EE, red) in addition to the downstream RNA hairpin. The EE form of the construct exhibits a diminished mKate intensity relative to the construct containing WT TM2, though it is still greater than the intensity associated with the expression of a construct containing silent mutations that disrupt the slip-site (SSKO). These observations demonstrate that the membrane integration efficiency of TM2 modifies the PRF efficiency when the RNA hairpin is present. B) A histogram depicts the distribution of mKate intensities for cells expressing PRF reporter constructs containing the upstream nascent chain with either the WT TM2 (blue) or a mutant form of TM2 bearing to non-native glutamate residues (EE, red), but lacking the downstream RNA hairpin. The mKate intensities associated with constructs containing the WT or EE versions of TM2 are indistinguishable from those of cells expressing a construct containing silent mutations that disrupt the slip-site (SSKO). These observations demonstrate that the membrane integration of TM2 only stimulates PRF in the presence of the downstream RNA hairpin.

### Supplemental Table 1. Experimentally Characterized Sindbis Virus Polyprotein Variants

| Variant | Sequence† | Predicted $\Delta G_{app}$ of TM2 (kcal/ mol)* | Membrane Integration Efficiency of TM2** | Distance from TM2 to Slip Site (Residues) | Mean Force (pN)*** |
|---|---|---|---|---|---|
| WT | LTPYALAPNAVIPTSLALLCCVRSANAE TFTETMSYLWSNSQPFFWVQLCIPLAAF IVLMRCCSCC | 1.9 | 0.44 ± 0.04 | 45 | 29.2 |
| EE | LTPYALAPNAEEPTSLALLCCVRSANAE TFTETMSYLWSNSQPFFWVQLCIPLAAF IVLMRCCSCC | 5.3 | 0.11 ± 0.03 | 45 | 26.9 |
| LL | LTPYALAPNAVIPLLLALLCCVRSANAE TFTETMSYLWSNSQPFFWVQLCIPLAAF IVLMRCCSCC | 0.2 | 0.51 ±0.04 | 45 | 33.3 |
| -15 | LTPYALAPNAVIPTSLALLCCVRSANAE TFWVQLCIPLAAFIVLMRCCSCC | 1.9 | 0.77 ± 0.03 | 30 | 21.3 |
| -10 | LTPYALAPNAVIPTSLALLCCVRSANAE TFTETMFWVQLCIPLAAFIVLMRCCSCC | 1.9 | 0.62 ± 0.04 | 35 | 26.8 |
| -5 | LTPYALAPNAVIPTSLALLCCVRSANAE TFTETMSYLWSFWVQLCIPLAAFIVLMR CCSCC | 1.9 | 0.50 ± 0.04 | 40 | 30.8 |
| +5 | LTPYALAPNAVIPTSLALLCCVRSANAE TFTETMSYLWSNSQPFGGGSGFWVQLC IPLAAFIVLMRCCSCC | 1.9 | 0.41 ± 0.04 | 50 | 25.4 |
| +10 | LTPYALAPNAVIPTSLALLCCVRSANAE TFTETMSYLWSNSQPFGGGSGGGGSGF WVQLCIPLAAFIVLMRCCSCC | 1.9 | 0.37 ± 0.04 | 55 | 25.4 |
| +15 | LTPYALAPNAVIPTSLALLCCVRSANAE TFTETMSYLWSNSQPFGGGSGGGGSGG GGSGFWVQLCIPLAAFIVLMRCCSCC | 1.9 | 0.35 ± 0.04 | 60 | 25.9 |

†The amino acid sequence from the N-terminal residue of TM2 (L725) through the C-terminal residue of TM3 (C790) is shown for each variant. The sequences of TMs 2 & 3 are shown in red and orange, respectively. Point mutations are shown in blue. Inserted residues are shown in green.
* Values reflect the minimum $\Delta G_{app}$ value as determined from a sequence scan of the full-length structural polyprotein using the $\Delta G$ predictor. (18)
** Values are derived from CGMD, and reflect the percentage of trajectories in which TM2 was found to adopt a transmembrane orientation.
*** Values are derived from CGMD, and reflect the average force on the nascent chain while the ribosomal slip site occupies the ribosomal P-site. The error associated with these values is ≤ 0.1 pN.

### Supplemental Table 2. Coarse Grained Molecular Dynamics Simulation Parameters.

| | σ | ε | $x_m$ | $x_r$ | $r$ |
|---|---|---|---|---|---|
| Open-LLL | 1 | 0.38 | -0.40 | 1.41 | 2.81 |
| Closed-LLL | 0.8 | 0.42 | 0.57 | 0.38 | 3.39 |
| Open-DDD | 0.91 | 0.66 | -0.23 | 3.61 | 1.83 |
| Closed-DDD | 1.13 | 0.45 | -0.06 | 2.05 | 2.31 |
| Open, Confined-LLL | 1.18 | 0.85 | -0.40 | 1.41 | 2.81 |
| Closed, Confined-LLL | 0.82 | 1.93 | 0.57 | 0.38 | 3.39 |
| Open, Confined-DDD | 1.13 | 0.30 | -0.23 | 3.61 | 1.83 |
| Closed, Confined-DDD | 1.19 | 0.68 | -0.06 | 2.05 | 2.31 |

The CGMD model parameters describing the interactions between the channel and the nascent chain have been updated using a Bayesian uncertainty quantification framework.[43] The parameterization here also addresses the systematic under-estimation of the integration probability found in earlier work by scaling the barrier of translocation for the DDD tri-peptide in the reference MARTINI PMF[37] before fitting, scaling the barrier by a factor of 0.5 was found to give the best fit with experiments.[27] The interaction between the channel and the nascent chain were parameterized explicitly for the most hydrophobic and hydrophilic bead types, composed of three leucines (LLL) and three aspartates (DDD), respectively. The interactions with other bead types are then determined by linear interpolation between these extremes. Separate parameters were obtained for interactions with the open and closed channel, as well as for the two bead types in the Sec translocon, which we call standard and confined. Which beads take on the confined parameters is determined by three parameters: $x_m$, which defines the start of the confined region, $x_r$ which defines the thickness of the region in the vertical direction, and $r$, which defines the radius of the region in the xy-plane. Each channel bead interacts with the nascent polypeptide through a 6-12 Lennard-Jones potential, described by the parameters σ and ε. In addition to the adjustment to the channel-nascent chain potential, the positioning of the lipid was also adjusted slightly. In particular, the lipid was defined to have a height of 1.99, and is excluded from a region with radius of 2.03 from the center of the channel. All units are in reduced (Lennard-Jones) units. Precise definitions of each parameter and interaction can be found in reference 37.
